# Low-dose ketamine tilts the cortical excitation–inhibition balance irrespective of systemic physiological response

**DOI:** 10.64898/2026.09.17.752361

**Authors:** Judith Kunze, Justus P. Student, Sarah Tune, Henrik Oster, Stefanie Schmidt, Harald Ihmsen, Alexander Tzabazis, Benedikt Lorenz, Carla Nau, Jonas Obleser

## Abstract

Non-invasive proxies of cortical excitation–inhibition (E:I) balance, are increasingly used to understand human neural dynamics; especially so the aperiodic (1/f) electroencephalographic (EEG) exponent. Ketamine, an NMDA-receptor antagonist, is thought to shift this balance toward excitation. Propofol, a GABA_A_-receptor agonist, should behave oppositely. Here we show that these pharmacological manipulations at low doses tilt the noninvasive read-out of E:I balance as predicted, and we demonstrate this effect to be robust against potential mediation by the ketamine-related systemic cardiovascular response. In a single-blind, placebo-controlled crossover study, 25 healthy adults received low, subanesthetic doses of ketamine, propofol, or placebo by target-controlled infusion during resting-state EEG. We further analyzed electrocardiogram (ECG) and dissociating ratings. Ketamine flattened the aperiodic exponent (i.e., a shift toward excitation) and raised heart rate and blood pressure; propofol exerted its effects in the opposite direction, while both agents reduced alpha oscillatory power. Critically, the cardiovascular response did not mediate the cortical E:I shift. Controlling for heart rate left the ketamine effect not only intact but numerically stronger, suggesting that E:I balance findings might even be underestimated when ignoring the systemic physiological response.

## Introduction

The balance of excitatory and inhibitory (E:I) signals within the brain is essential for efficient and stable neural functioning [1,2]. Disturbances of this balance have been associated with various neuropsychiatric disorders, including schizophrenia [3] and autism spectrum disorder [4].

Measuring E:I balance in humans is feasible using non-invasive electroencephalography (EEG). The aperiodic (1/f) spectral exponent of the EEG power spectrum has emerged as a proxy for E:I balance [5]: Flatter 1/f slopes are associated with greater excitation, and steeper slopes with greater inhibition [5]. The E:I balance is not a static trait, but can be state-dependent. For example, E:I balance shifts under sleep deprivation (hyper-excitation) and postprandial somnolence (hypo-excitation) [6]. Additionally, E:I balance changes systematically throughout the day [7] as well as by pharmacological manipulations [8,9]. Ketamine blocks N-methyl-D-aspartate (NMDA) receptors, inducing NMDA-receptor hypofunction in gamma-aminobutyric acid (GABA)-ergic inhibitory interneurons [10] and thereby shifting the E:I balance toward excitation [11] even at low, subanesthetic doses [12]. In contrast to ketamine, hypnotic doses of propofol have been shown to shift E:I balance toward inhibition by enhancing GABA_A_-receptor activity [11]. In addition to their impact on E:I balance, both substances also affect alpha power. While hypnotic dosage of propofol leads to increased frontal alpha power [13,14], the effect at subanesthetic doses is less known. Ketamine suppresses alpha power at both anesthetic and subanesthetic doses [15]. Ketamine’s, but not propofol’s, effect at low-dose on dissociative experience is well known [16,17].

Ketamine and propofol do not have an isolated effect on the brain, they additionally affect cardiovascular function. Subanesthetic doses of ketamine, for example, raise blood pressure and heart rate [18,19]. In contrast, propofol administration lowers blood pressure through peripheral vasodilation, and can cause bradycardia [20]. When making assumptions about central E:I balance, one has to bear in mind that both drugs have effects on the cardiovascular system. Over the past years it has become clear that neural activity can be shaped and influenced by body rhythms [21]. For example, a coupling between cardiac and neural rhythms has been demonstrated at rest [22]. Furthermore near-threshold electrical stimuli are detected more often during diastoles than during systoles [23,24], accompanied by changes in perceptual sensitivity and by suppression of late somatosensory-evoked potential components [23]. The neural response to the heartbeat itself also matters, as larger heartbeat-evoked potentials precede lower detection rates and a more conservative decision-making criterion [23].

Beyond such brain-body coupling, cardiac activity can also “contaminate” neural recordings. Schmidt et al. [25] showed that the aperiodic component recorded at surface sensors originates from multiple physiological sources, rather than from cortical activity alone, and that common artefact-rejection approaches such as independent component analysis may not be sufficient to separate cardiac from neural contributions. In a pharmacological within-subject design this concern becomes acute: since ketamine and propofol shift heart rate and blood pressure in opposite directions, any drug-induced change in the aperiodic exponent could in principle reflect altered cardiac contributions rather than a genuine cortical shift in E:I balance.

To date, however, drug-induced changes in the aperiodic exponent and drug-induced changes in cardiovascular function have largely been studied in isolation. It therefore remains unclear to what extent pharmacologically induced shifts in the aperiodic exponent reflect cortical E:I balance, concurrent cardiac changes, or a combination of both.

In the current study, we set out to close this gap by testing the following hypotheses (Fig. 1a). First, we expected both substances to alter cardiovascular function, as described above in opposite directions. Second, we expected the pharmacological manipulation to influence cortical alpha power (green arrow) and shift the E:I balance, proxied by the aperiodic spectral exponent, in opposite directions: with ketamine pushing toward excitation and propofol toward inhibition, proxied by the aperiodic spectral exponent. Additionally, we tested whether drug-induced changes in cardiovascular function (i.e. heart rate) mediate the effect of drug condition on the E:I balance (black arrows). We further examined how ketamine administration, E:I balance, and cardiovascular responses jointly or separately contribute to subjective dissociative experiences (pink arrows).

**Figure 1:**
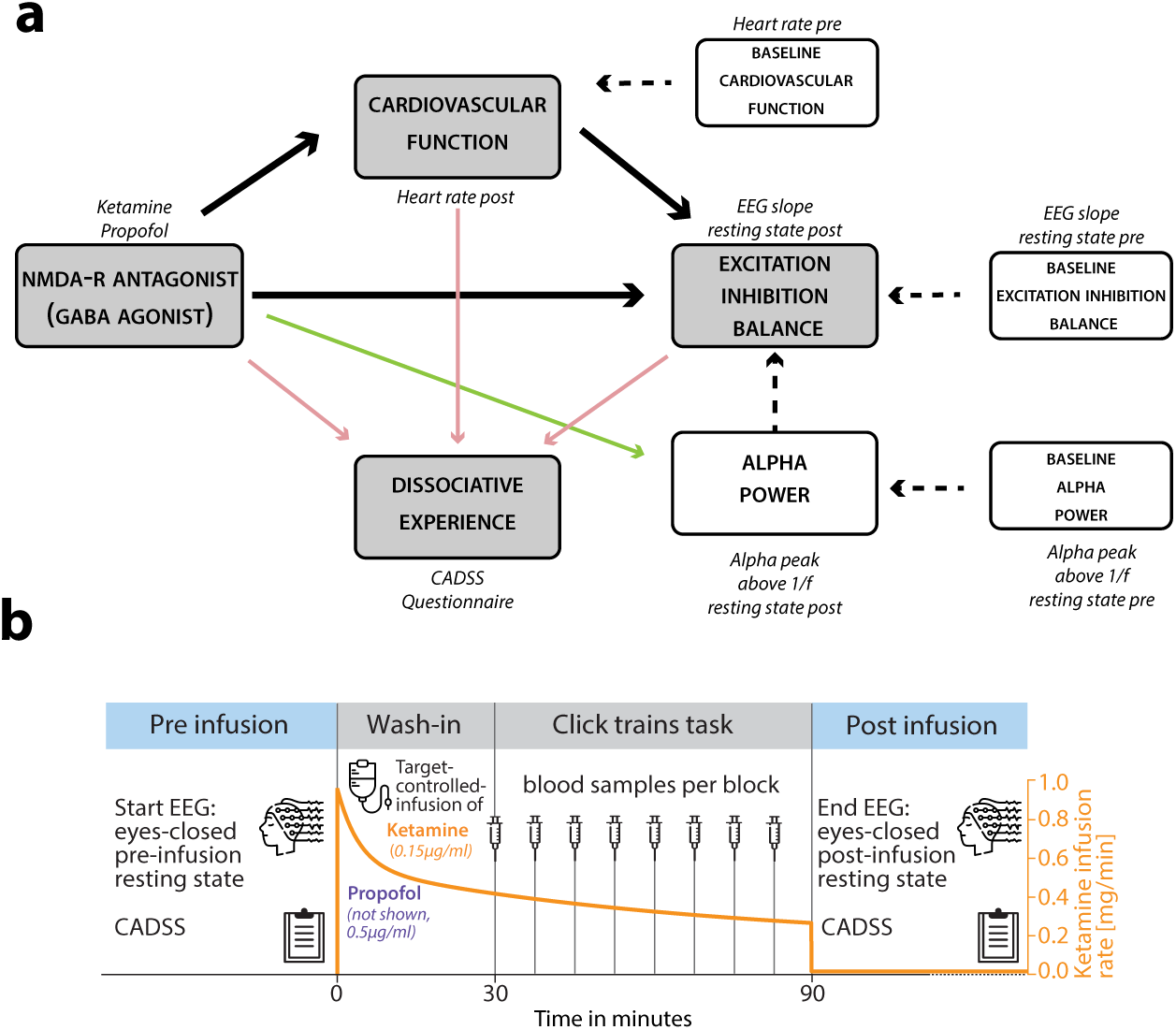
Overview hypothesis and study design. **(a)** Hypothesized relationships between latent (grey) and manifest (white) variables; italic text indicates the corresponding measure. Solid arrows are hypothesized directed effects, with colors representing different hypotheses. Black: effects of NMDA antagonism/GABA agonism on cardiovascular function and on E:I balance with cardiovascular function tested as a potential mediator of the drug effect on E:I balance; Green: drug effect on alpha power; Pink: predictors of dissociative experience. Dashed lines are potential covariates (pre-drug baselines, and alpha power on the E:I estimate). **(b)** A practice block with pre-infusion resting-state EEG and CADSS was followed by target-controlled infusion of ketamine (target plasma concentration 0.15 µg/ml) or propofol (0.5 µg/ml, infusion profile not shown); the orange line gives an example of ketamine infusion rate (right axis). After a wash-in rest period, participants performed the click-trains task (data not shown), with one blood sample drawn per block. The task ended after approximately 90 min, followed by a post-infusion CADSS and resting-state EEG recording.

## Methods

### Participants

The experiment was conducted on a sample of 32 healthy participants. Seven of them did not complete all conditions. Therefore, the final analysis population was 25 participants (15 female). The participants were predominantly students at the University of Lübeck, with a mean age of 23.56 years (SD = 4.54), with no psychological or physical diseases and normal hearing. Prior to the first experimental session, the participants attended an informational meeting with the designated physician, and provided their written consent. Procedures were approved by the ethics committee of the University of Lübeck and were in accordance with the Declaration of Helsinki. Participant characteristics are shown in Table S1.

### Study design and study plan

A randomized, single-blinded, placebo-controlled, three-condition crossover study was applied including ketamine, propofol, and placebo condition.

Participants were recruited via the online recruitment system for economic experiments [26]. Each participant completed three EEG sessions on three separate days (minimum between-session interval of 7 days), each including an auditory decision-making task [27]. Task data, EEG and behavioral results, are not part of this analysis. For safety, all sessions took place in the intensive care unit at the University Hospital Schleswig-Holstein, Campus Lübeck under continuous monitoring by a physician.

While not the main focus of the current investigation, in every placebo session we determined the individual dim-light melatonin onset (DLMO) as a marker of internal circadian phase. Because direct assessment of DLMO requires repeated sampling under controlled dim-light conditions and is therefore time-consuming and burdensome for participants, we instead used the predicted DLMO derived from blood cell gene expression profiles [28]. This approach infers internal circadian time from the transcriptional profile of peripheral blood mononuclear cells (PBMCs) of a single blood sample and has been validated against laboratory-assessed DLMO [28]. The participants were instructed to refrain from eating for six hours and drinking for two hours before each session. Approximately 24 hours after each EEG session, participants underwent an MRI scan lasting about 1.5 hours. The MRI results are not shown here. Participants received €20 per hour as monetary compensation.

Within each EEG session, procedures followed a fixed sequence (Fig. 1b). The session began with an eyes-open and closed resting-state EEG (4 minutes each, pre) and an initial Clinician-Administered Dissociative States Scale (CADSS) [29] assessment. After a brief training period, oxygen (2 L/min) was administered via nasal cannula, and the intravenous infusion was started. We allowed for a 30-minute wash-in period to ensure a steady-state concentration throughout the body [30]. Participants then completed an auditory decision-making task, which lasted approximately 70 minutes. Immediately after task completion, the infusion was stopped, and a second eyes-open and -closed resting-state EEG was recorded (4 minutes each, post), and the CADSS was administered another time. In total, each session took around 4 hours.

### Target-controlled infusion

The application of both intervention agents was conducted using a target-controlled infusion (TCI) pump (Arcomed AG; Kloten, Zurich, Switzerland) to maintain a constant drug plasma levels during the measurement. The targeted ketamine plasma level was 0.15 µg/mL estimated based on the Domino model [31]. In comparable studies involving healthy adults, it has been shown that neuropsychological effects occurred even at this low-dose level [8,32,33]. For the administration of propofol, the Schnider model was used, which adjusts the infusion dose to the subject based on both weight and age [34,35]. The targeted plasma concentration was set to 0.5 µg/mL. In other studies, antiemetic effects were already observed at comparable plasma levels without affecting the state of consciousness of the subjects studied [36–39].

### Plasma concentrations

In total eight blood samples were collected from an intravenous cannula within each condition to quantify the accuracy of the TCI. The first blood sample was taken 30 minutes after starting the infusion, assuming an intracerebral steady state conditions had been reached by then [30]. The remaining seven blood samples were subsequently drawn in intervals of 8 minutes coinciding with auditory task block breaks.

For both agents, sample preparation was performed by protein precipitation using methanol. Quantification of propofol and ketamine concentrations in human plasma was carried out using ultra-high-performance liquid chromatography coupled with tandem mass spectrometry. Detection was conducted in positive electrospray ionization mode with deuterated internal standards as reference compounds. The lower limits of quantification were 50 ng/ml for propofol and 20 ng/ml for ketamine, respectively. The inter- and intra-assay errors were within ±5 % for propofol and within ±10% for ketamine, respectively. Additional methodological details have been described previously [40,41].

### EEG measurement and preprocessing

The EEG was recorded from 64 electrodes (ActiCap/ActiChamp; Brain Products, Gilching, Germany) at a sampling rate of 1,000Hz and the impedances were kept below 20 kΩ with online reference TP9.

EEG preprocessing was conducted with a custom multi-stage pipeline using EEGLAB (v2024.1) and FieldTrip toolboxes in MATLAB (The MathWorks Inc., Natick, USA) [27]. Independent component analysis (ICA) was performed using the runica algorithm. Components were automatically classified using the ICLabel plugin [42]. Components exceeding 90% probability of representing non-brain sources (e.g., eye, muscle, heart, line noise, channel noise, or other) were removed before importing into FieldTrip for further preprocessing and analysis. Channels with impedances >50 kΩ based on the EEG amplifier log files or considered as noisy by visual inspection were identified and interpolated using spherical spline interpolation. EEG data were re-referenced to average and demeaned. Eyes-closed resting state recordings were cut to a uniform length of the last 3 recorded minutes each. The data were broadband filtered using custom-designed FIR filters (high-pass: 0.3 Hz) and resampled to 250 Hz.

### Cardiovascular monitoring

Cardiovascular monitoring, including continuous electrocardiogram (ECG), heart rate, and intermittent blood pressure measurement with an interval of three minutes, was initiated 20 minutes before the start of the infusion. The mean arterial pressure and heart rate were further analyzed from the processed data.

We recorded an additional ECG (sampling rate 1,000Hz) alongside the EEG to optimize synchronization between both signals. Heart rate was derived from a single-lead ECG channel and trimmed to the corresponding eyes-closed resting-state segment. The ECG signal was processed using the default NeuroKit2 [43] cleaning method, and R-peaks were detected with the NeuroKit2 algorithm. Subsequently, detected R-peaks were corrected using an iterative artifact-correction procedure and thereafter the instantaneous heart rate was calculated from the resulting inter-beat (RR) intervals as 60000/RR (ms) as well as the mean heart rate. After visual inspection five recordings were marked as noisy and were replaced by initial ECG which was conceptually used for monitoring during the measurement.

### Calculation of alpha power

Spectral analysis and aperiodic fitting were done using Python 3, the scipy.signal.welch function and the FOOOF toolbox [44], respectively.

Power spectra were computed using Welch’s method (Hann window, segment length = 1024 samples (∼0.244 Hz), 50% overlap), separately for each channel and then averaged across all channels.

FOOOF models were fit to the power spectra (1–50 Hz) with the following parameters: peak width limits of 1–8 Hz, maximum number of peaks = 8, minimum peak height = 0.05, peak threshold = 2.0 SD. Models were fit using fixed aperiodic mode.

To isolate oscillatory alpha power from the aperiodic (1/f) background, the aperiodic component of the FOOOF fit and the full model fit were first converted from log-power to linear power, and the aperiodic fit was then subtracted from the full model fit in linear space, yielding a 1/f-corrected (periodic) power spectrum. Any residual negative values arising from this subtraction were set to zero. Alpha power was computed as the area under this corrected spectrum within the 8–12 Hz range using trapezoidal numerical integration, but only if FOOOF identified a discrete spectral peak with a center frequency within this range and a bandwidth not exceeding the width of the band (BW ≤ 4 Hz). This criterion was applied to ensure that the resulting estimate reflects a genuine narrowband alpha oscillation rather than spectral energy attributable to a broader, non-band-limited periodic component overlapping the alpha range.

### Calculation of the aperiodic exponent as a proxy for E:I balance

The aforementioned power spectra computation was used and aperiodic (1/f) spectral parameters were fit to 30–50 Hz range [5]. Model settings were: peak width limits: 1–8 Hz, maximum number of peaks = 8, minimum peak height = 0.05, peak threshold = 2.0 SD. From each fit we extracted the exponent as a proxy for E:I balance.

### Measuring dissociative experience

To capture the subjective dissociative feeling, we asked the participants at the beginning and another time at the end of the experiment the questions from the CADSS, where participants answer on a 5 point Likert scale (higher scores indicate more dissociative experience). We used the simplified 6-item modification in the validated German version [45], which was specifically constructed for monitoring dissociative effects of subanesthetic ketamine infusions [46].

### Statistical analysis

All continuous variables were z-standardized prior to analysis and condition (ketamine, placebo, propofol) as a factor was simple-effect coded. Linear mixed-effects models were fit in R Statistical Software (Version 4.3.2 R Foundation for Statistical Computing, Vienna, Austria) using *lme4* and *lmerTest,* with restricted maximum likelihood (REML) estimation, Satterthwaite-approximated degrees of freedom, and participant-specific random intercepts account for repeated measurements within subjects.

Each post-infusion outcome (alpha power, heart rate, aperiodic exponent, CADSS) was regressed on its respective pre-infusion baseline value, condition, and session order.

To test whether the effect of ketamine on the aperiodic exponent was mediated by concurrent changes in heart rate, we conducted a causal mediation analysis using the quasi-Bayesian Monte Carlo approximation implemented in the R package *mediation* [47]. The analysis was restricted to the ketamine and placebo sessions, with the condition entered as a binary treatment (0 = placebo, 1 = ketamine). The mediator and outcome models corresponded to the heart rate and aperiodic exponent models described above, with post-infusion heart rate added as an additional predictor in the outcome model. The average causal mediation effect (ACME), average direct effect (ADE), and total effect were estimated from 5,000 Monte Carlo draws, with 95% confidence intervals and two-sided *p* values derived from the resulting quasi-Bayesian distributions.

All models and full table results are shown in Table S2–S11.

## Results

### Plasma level concentrations hit targeted levels

The mean ketamine plasma concentration across all participants and all eight samples was 138.21 (SD = 29.02) ng/mL. Initially, the average plasma level undershot the target level of 150 ng/mL and subsequently increased over time. During the final 30 minutes of the observation period, a steady state was reached at around 160 ng/mL. Within the propofol session, the mean propofol plasma level concentration was 486.32 (SD = 118.31) ng/mL. At first, the average plasma level was slightly below the target level and increased throughout the observation period. An approximate steady state of around 550 ng/mL was also reached also within the last 30 minutes. Notably, two participants showed markedly elevated and highly fluctuating propofol concentrations. The trajectories of the plasma level concentration of both conditions are shown in figure 2a.

**Figure 2:**
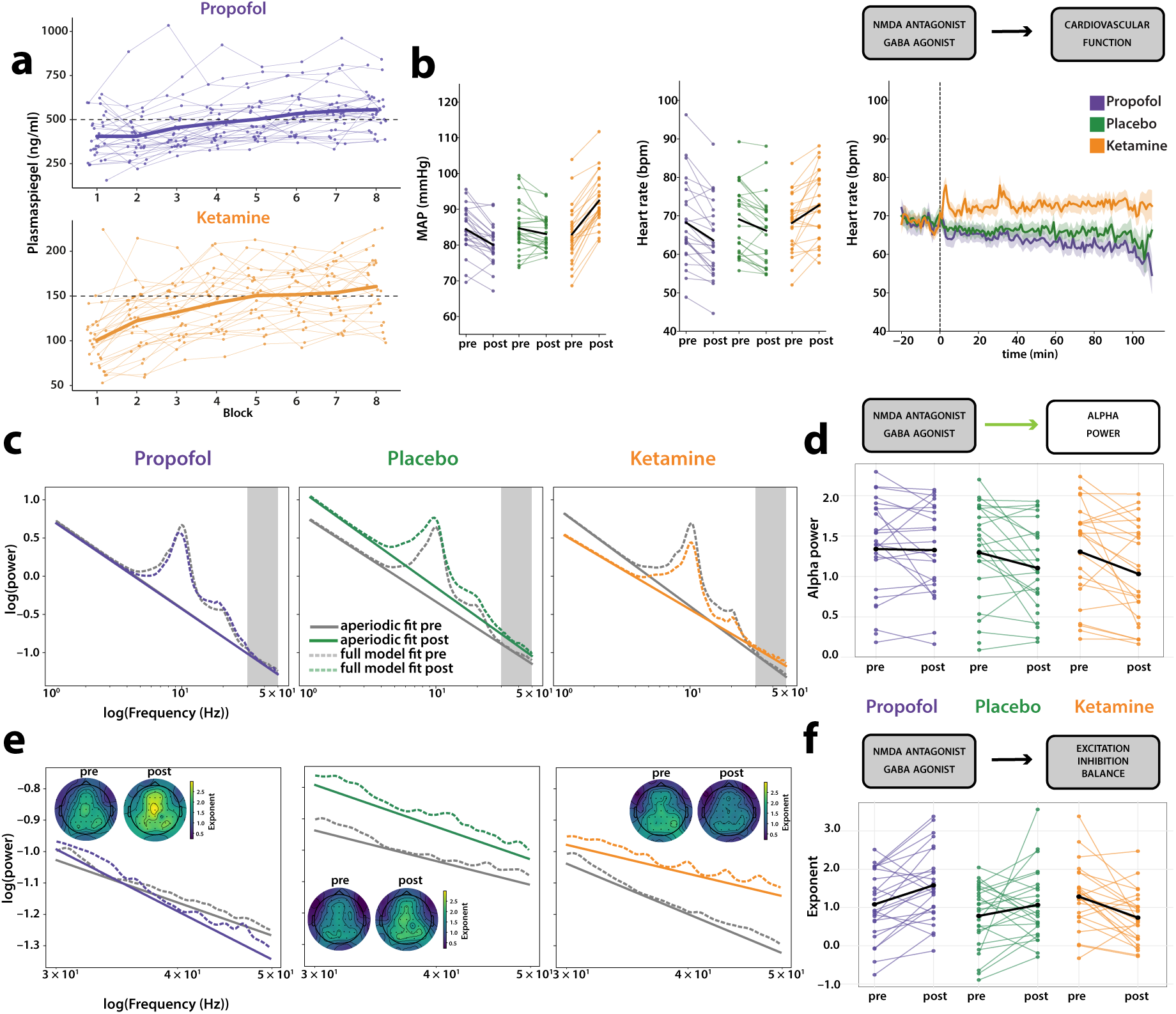
Pharmacokinetics, cardiovascular response and resting-state EEG spectral changes under propofol, placebo and ketamine. Purple, propofol; green, placebo; orange, ketamine. Thin lines show individual participants, thick black lines the group mean. Grey boxes state the hypothesis addressed in the adjacent panel. **(a)** Plasma concentrations across the experimental blocks for propofol (top) and ketamine (bottom); dashed line, targeted concentration. **(b)** Mean arterial pressure (MAP) (left) and heart rate (HR) (middle) pre to post infusion, and the continuous heart-rate time course relative to infusion onset. Ketamine increased both MAP and HR, whereas both measures decreased under placebo and, more markedly, under propofol. **(c)** Grand-average resting-state power spectra (log–log, 1–50 Hz), averaged across channels and participants. Grey, pre-infusion; colored, post-infusion; solid lines, aperiodic component of the FOOOF fixed-mode fit; dashed lines, full model fit. The grey shaded area marks the 30–50 Hz range shown in (e). **(d)** Aperiodic-adjusted alpha power (8–12 Hz periodic component of the FOOOF fit) pre to post, per condition from the spectrum in (c). Thin lines show individual participants, the black line the group mean. For visualization, the y-axis is truncated at 70.; one participant (participant 25) exceeds this range in the ketamine (130; 90) and propofol (131; 69) conditions and is labelled accordingly. This participant was retained in all analyses and contributes to the group means shown. Participants without an identifiable alpha peak at one or both timepoints (n = 3 ketamine, 4 placebo, 3 propofol) are not plotted. **(e)** Grand-average spectra over the 30–50 Hz fitting range, plotted as in (c). Insets, scalp topographies of the channel-wise 30–50 Hz aperiodic exponent pre and post infusion. **(f)** Aperiodic exponent estimated over 30–50 Hz, pre to post, per condition. Ketamine flattened the spectrum, i.e. the exponent decreased, whereas it increased slightly under placebo and propofol.

### Ketamine increased blood pressure and heart rate whereas propofol shows opposite effects

Within the ketamine session, the baseline mean arterial pressure was 83.0 (*SD* = 7.9) mmHg. During infusion the mean arterial pressure increased to 92.3 (*SD* = 7.0) mmHg. By contrast, propofol decreased arterial pressure from 84.4 (*SD* = 6.3) mmHg to 80.0 (*SD* = 5.3) mmHg. In the placebo control session, we observed prior to infusion a mean arterial pressure of 84.7 (*SD* = 6.8) mmHg versus 83.1 (*SD* = 4.7) mmHg post infusion. In statistical analysis, post-infusion mean arterial pressure differed significantly by condition (*F*(2, 48.88) = 110.21, *p* < .001), adjusting for pre-infusion values. Compared to placebo, mean arterial pressure was markedly higher under ketamine (*β* = 1.36, *SE* = 0.12, *t*(34.3) = 11.42, *p* < .001) and significantly lower under propofol (*β* = −0.36, *SE* = 0.12, *t*(34.7) = −2.93, *p* = .006; Fig. 2b).

Ketamine infusion increased the heart rate from 68.1 (SD = 7.2) bpm to 72.8 (SD = 8.6) bpm. Within the propofol session, there was a decrease from 68.2 (SD = 10.7) bpm to 63.6 (SD = 9.8) bpm. Further, after placebo infusion the heart rate dropped from 69.1 (SD = 8.9) bpm to 66.1 (SD = 8.9) bpm. Post-infusion heart rate differed significantly by condition (F(2, 38.08) = 42.61, p < .001), controlling for pre-infusion heart rate. Heart rate was significantly higher under ketamine infusion (β = 0.76, SE = 0.11, t(38.07) = 7.072, p < .001), while the propofol condition did not differ significantly from the placebo condition (β = −0.17, SE = 0.11, t(38.44) = −1.55, p = .129) (Fig. 2b).

### Alpha power decreased under ketamine and propofol

The analysis of alpha power revealed a significant modulation by drug condition (*F*(2, 42.5) = 8.41, *p* < .001). Relative to the placebo condition, we observed reduced oscillatory alpha power following both ketamine (β = −0.51, *SE* = 0.12, *t*(42.3) = −4.10, *p* < .001) and propofol infusion (β = −0.25, *SE* = 0.12, *t*(42.4) = −2.06, *p* = .046; Fig. 2c and 2d, S1).

### Low doses of ketamine and propofol shift E:I balance as expected by their pharmacological response profile

Aperiodic activity was parameterized with the FOOOF algorithm which captured the observed spectra well (Mean R^2^ = .714). Fitted full-model and aperiodic spectra together with the scalp distribution of the exponents are shown in Fig. 2e.

First we established that the approach of estimating aperiodic slope (i.e., the exponent) from three minutes of resting-state EEG is reliable and yields a stable, trait-like signature within individuals. Indeed, this was the case: pairwise Pearson correlations of the pre-infusion exponent between sessions (i.e., placebo, ketamine, propofol) yielded satisfactory levels of reliability (all r [.66; .74], all t(23) > 4.15, all p < .001).

Second, pharmacological treatment altered the exponent as expected (*F*(2, 70) = 9.28, *p* < .001). Relative to placebo, ketamine reduced the exponent (*b* = −0.62, *SE* = 0.24, *t*(70) = −2.55, *p* = .013), that is, it flattened the aperiodic slope, consistent with a shift of the excitation–inhibition balance towards excitation. Propofol showed a trendwise increase in the exponent (*b* = 0.40, *SE* = 0.24, *t*(70) = 1.68, *p* = .09; (Fig. 2f)).

Third, testing the hypothesis of opposing effect of ketamine versus propofol on the exponent post most directly, a model using a contrast of ketamine > propofol and nulling the impact of placebo yielded an average shift of *b* = –1.027 standard deviations in exponent, *SE* = 0.24, *t*(70) = –4.289, *p* < 0.001.

### Effects of ketamine on E:I balance were robust under cardiovascular control

Because ketamine reliably increased both heart rate and the flattening of the aperiodic slope, the change in the exponent could in principle be a consequence of the cardiovascular response rather than a genuine change in cortical excitation–inhibition balance. We therefore decomposed the total effect of ketamine on the aperiodic exponent into an indirect path through post-infusion heart rate and a direct path, using causal mediation analysis. The total effect of ketamine on the aperiodic exponent was significant (−0.62, 95% CI [−1.17, −0.05], *p* = .032) and was carried by the average direct effect (ADE = −0.79, 95% CI [−1.41, −0.20], *p* = .012). The average causal mediation effect was not significant and, notably, of opposite sign to the total effect (ACME = 0.17, 95% CI [−0.06, 0.44], *p* = .167), indicating that there is no evidence that the ketamine-induced flattening of the aperiodic slope is transmitted through the cardiovascular response.

Rather than accounting for the drug effect, adjusting for the cardiovascular response made it larger (Fig. 3). Without heart rate in the model, ketamine reduced the post-infusion exponent by *β* = −0.62 (*SE* = 0.28, *t*(45) = −2.22, *p* = .032). With post-infusion heart rate included, the estimate increased in magnitude to *β* = −0.79 (*SE* = 0.31, *t*(44) = −2.60, *p* = .013). The difference between the two ketamine estimates is exactly the indirect path through heart rate, i.e. the ACME reported above (−0.62 − (−0.79) = 0.17).

**Figure 3:**
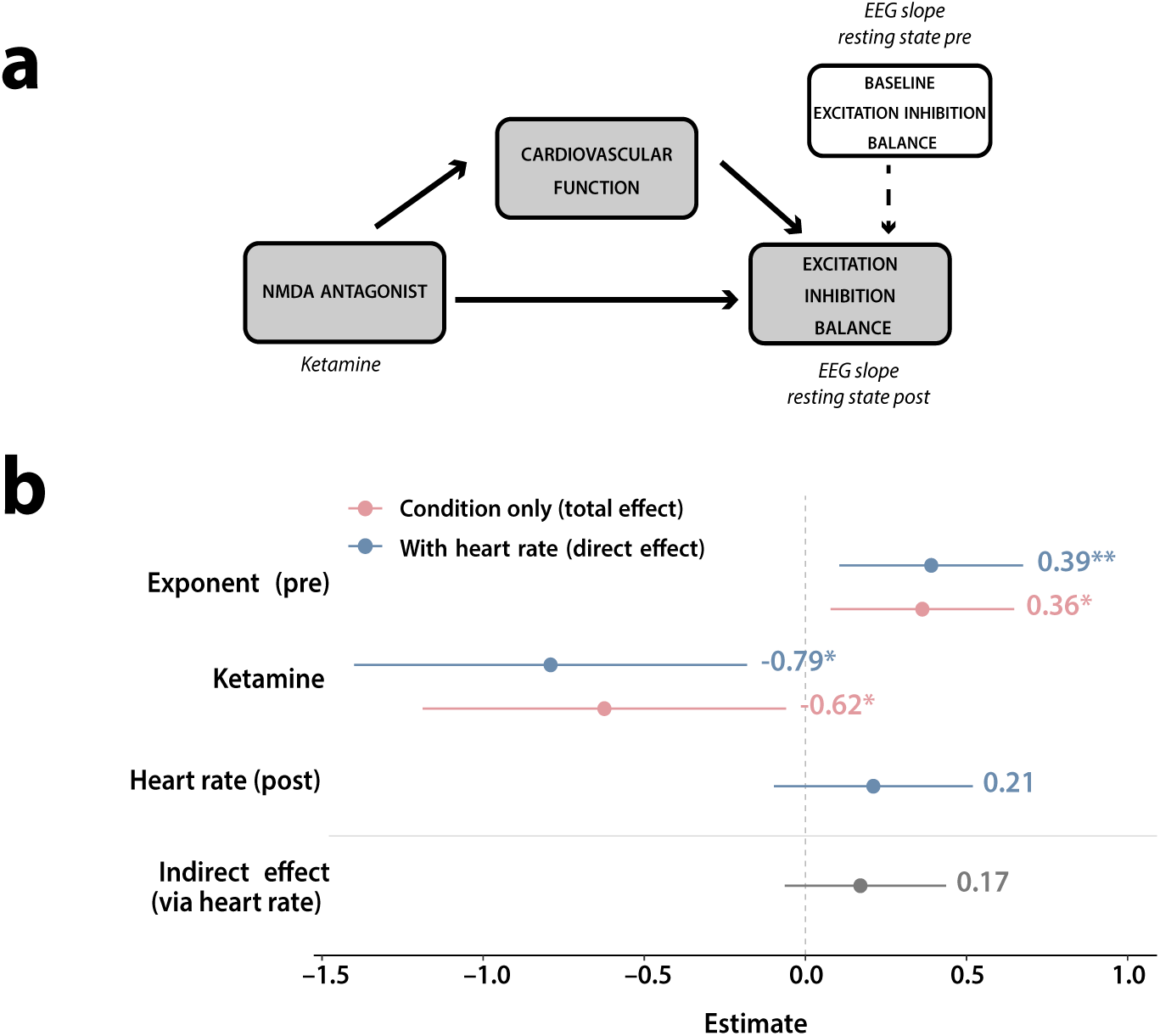
Mediation analysis of the ketamine effect on the aperiodic exponent by the cardiovascular response. **(a)** Hypothesized paths: **(b)** Model estimates for the post-infusion 30–50 Hz aperiodic exponent. Light pink, condition-only model (total effect of ketamine relative to placebo, adjusted for the baseline exponent); Light blue, the same model with post-infusion heart rate added as a covariate (direct effect); grey, the indirect effect via heart rate. Points show standardized regression estimates, horizontal lines the 95% confidence interval; the dashed vertical line marks zero. \**p* < 0.05, \*\**p* < 0.01. It implies that the effect of ketamine on E:I balance is not mediated, maybe even suppressed by heart rate.

### The ketamine-induced dissociative experience is not explained further by cardiovascular function or E:I balance

Ketamine infusion produced a robust increase in dissociative symptoms (*β* = 1.53, *SE* = 0.23, *t*(23.91) = 6.51, p < .001). Neither the post-infusion aperiodic exponent (*p* = .433) nor heart rate (p = .158) explained additional variance in CADSS scores beyond the condition effect. Additional mediation analysis found no evidence that either physiological measure mediates the ketamine effect on dissociation: the ACME was near zero and non-significant for both 1/f exponent (ACME = –0.174, 95% *CI* [–0.61, 0.25], *p* = .42) and heart rate (ACME = –0.315, 95% *CI* [–0.78, 0.09], *p* = .14), while the direct effect (ADE) remained large and significant in both cases (p < .001). The model comparison confirmed the pattern from the individual models (Fig. 4).

**Figure 4:**
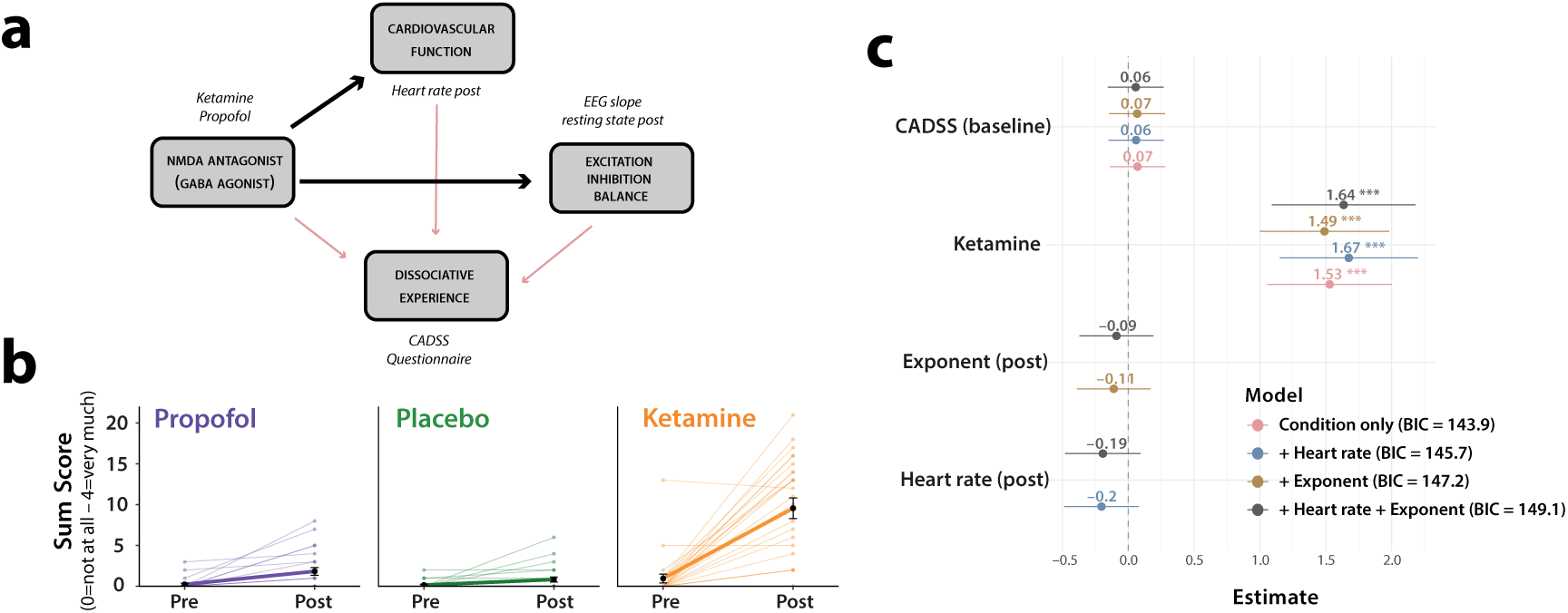
Joint effect of NMDA antagonist, cardiovascular function and E:I balance on dissociative experience. **(a)** Hypothesized paths: the NMDA antagonist (ketamine) may drive dissociative experience directly, or indirectly via cardiovascular function (post-infusion heart rate) or via E:I balance (post-infusion EEG slope); red arrows are the tested paths. **(b)** CADSS sum score (items rated 0 = not at all to 4 = very much) pre to post, per condition. Thin lines show individual sessions, thick lines the group mean. Dissociative symptoms increased steeply under ketamine, whereas most propofol and placebo sessions remained at or close to zero. **(c)** Estimates for the post-infusion CADSS score from four nested models, each adjusted for baseline CADSS: condition only (light pink), plus post-infusion heart rate (light blue), plus the 30–50 Hz aperiodic exponent (gold), and plus both (grey). Points show regression estimates, horizontal lines the 95% confidence interval; the dashed vertical line marks zero and the legend gives each model’s BIC. The ketamine estimate was large and essentially unchanged across models, while neither heart rate nor the aperiodic exponent explained additional variance; BIC favored the condition-only model. *** *p* < 0.001.

Note that our main results are all based, for stringency, on data from the respective eyes-closed sections of the resting-state EEG recordings (Figure 1). We repeated all main statistical models on the merged data of eyes closed and eyes open, with an additional nuisance regressor “eyes closed – eyes open”. The ketamine-induced decrease in the aperiodic exponent was preserved (b = −0.47, SE = 0.16, t(120) = −2.98, p = .003). Under propofol, the estimate showed the same positive trend as in the main analysis but qualitatively unchanged (b = 0.29, SE = 0.16, t(119) = 1.86, p = .07). Adding post-infusion heart rate as a further covariate again increased ketamine’s estimate slightly (b = −0.61, SE = 0.17, t(140) = −3.56, p < .001). It is therefore safe to conclude that our main findings — drug-dependence of the noninvasive read-out of E:I balance; independence of this effect from the drug-related cardiovascular response — derive independently from eye closure.

## Discussion

In this study we have shown that low, subanesthetic doses of ketamine and propofol shifted the aperiodic exponent broadly as their pharmacology predicts, lending support to scalp-level readouts of the spectral slope as indices of cortical E:I balance. Both drugs also influence cardiovascular components, especially heart rate, as expected: ketamine exerted a sympathoexcitatory cardiovascular effect and propofol a sympathoinhibitory effect. Crucially, ketamine’s effect on the E:I balance is not explained by the concomitant increase in cardiovascular activity.

First, target-controlled infusion allowed us to maintain this state across an experimentally relevant window, establishing a protocol we saw as worth exploring in future work. Further, our data showed that ketamine infusion even in subanesthetic doses could increase cardiovascular response reflected by a higher heart rate and elevated mean arterial pressure. This is in line with previous studies where other groups linked subanesthetic doses of ketamine use with higher values of blood pressure and heart rate in psychiatric patients [48] and in healthy young adults [49]. By contrast, propofol infusion decreased the mean arterial pressure and showed no statistically significant effect on heart rate [50] which was also reflected in our data. Notably, all post-measurement were obtained after infusion offset, when plasma concentrations had already begun to decline; the reported effects may therefore be underestimated.

Second, alpha power was reduced under both drugs relative to placebo. Because alpha power was estimated over and above the aperiodic 1/f component, this finding is not simply an epiphenomenon of the change in 1/f shape. It is notable that the two agents, namely ketamine as an NMDA-receptor antagonist and propofol as GABA_A_-receptor agonist, did not differ from one another on this measure. This result should nevertheless be interpreted with caution, as alpha power was calculated over the whole scalp. A spatially more specific analysis is warranted, particularly for propofol, where alpha power has been reported to increase over frontal regions at hypnotic doses [13]. Averaging across the whole scalp could therefore combine opposing local effects and mask topographically specific differences between conditions.

Third, the two agents moved the exponent in opposite directions: an NMDAR antagonist acting on inhibitory interneurons to shift the balance toward excitation, and a GABA_A_ agonist toward inhibition. While the ketamine effect was reliable the propofol shift consistent in direction though trend-level.

Fourth, we did our analyses under close measurement of potential autonomic-arousal responses, treating cardiac activity — heart rate, alongside blood pressure — as a candidate mediator. A mediation model was warranted here precisely because both drugs act strongly on the cardiovascular system. This analysis asks whether the drug’s effect on the aperiodic exponent is direct or is mediated through cardiovascular response. Yet controlling for the cardiovascular response did not diminish the drugs’ effect on the aperiodic exponent; if anything, the estimate increased, a pattern consistent with suppression rather than mediation — the pattern other studies may encounter when estimating cortical E:I changes under a drug that is also peripherally active. In short, statistically adjusting for cardiovascular response stabilized rather than neutralized the cortical E:I readout. Our finding of a possible suppression of the drugs’ direct effect when cardiovascular changes are left unmodelled is informative in several respects. First, it underscores the need for peripheral control and co-registration when working with centrally and peripherally active drugs [51] — a well-known, but often overlooked point. It is especially pertinent as neuroscience increasingly integrates central, autonomic, and bodily processes into its account of cortical function [52,53], with cardiac [54] and gastric [55] signals now shown to shape cortical dynamics directly.

Moreover, there have been important calls for caution in interpreting the Magneto- or electroencephalographic spectral slope readily as cortical E:I change. Schmidt et al. [25] raised the possibility that unaccounted or residual cardiac artefacts could account for apparent cortical changes. In the present data, however, the result point, if anything, the other way: heart rate was related to the exponent in a direction that opposed the drug effect, such that unmodelled cardiac influence would suppress rather than inflate the expected result. Accounting for it therefore recovered, rather than explained away, the cortical signal (Fig. 3).

In how far do our results hinge on specific methodological choices in EEG analysis? First, note that we intendedly used the 30–50 Hz spectra. The fitting range is not a trivial choice: a range dependence has been reported previously, where ketamine flattened the exponent in a high-frequency band while steepening it at low frequencies, so that over a broad range the opposing changes cancelled and ketamine became indistinguishable from wakefulness [56]. Additionally, fitting sub-ranges selected to avoid oscillatory peaks and knee frequencies, rather than a single broadband range, is established practice for this reason [57]. In our data, ketamine’s effect was robust to it, as the exponent also flattened over 1–50 Hz (Fig. S2), whereas propofol showed no change in the broadband fit. We decided for the 30–50 Hz range in which the E:I balance interpretation was originally derived. It lies above the oscillatory peaks and below the frequencies contaminated by spiking activity [5], and here the model’s link between E:I balance and spectral slope holds. In this vein, we also aimed to restrict the fit to a range containing no oscillatory peaks and thus avoiding any dependence of the aperiodic estimate on the concurrent fitting of periodic components. This cannot be achieved when the fitting range spans the alpha band. Also, we are aware that muscle activity can heavily influence the EEG in this frequency range [15]. We addressed this with ICA-based artefact rejection [58]. Notably, the number of muscle-labelled ICA components was highest under propofol (the condition which the exponent steepened, whereas broadband muscle activity would flatten it). The reported cortical pattern of drug effects is therefore dissociated from the degree of myogenic contamination.

Second, a remaining question is one of potential drug-related temporal dynamics, which our analyses might not have captured. As with alpha power, exponents were averaged across all electrodes, and potential topographical differences are not subject of this study. Exponents were likewise averaged across the full 3-min resting-state segment and thus treated as a static property of each condition. This is a simplification, as the aperiodic exponent is itself read as a frequency-domain index of neural variability, and variability of this kind is not stationary but fluctuates within an individual on the timescale of seconds to minutes [59]. Each spectral estimate is an aggregate over pharmacological states that need not be constant: drug effects on cortical activity can reverse in sign over tens of minutes, as when psilocybin initially increases sound-evoked response amplitude in mouse auditory cortex but decreases it roughly 30 min post-dose, while shared (noise-correlated) variability rises [60]. Time-resolved analyses within and across segments are a natural next step, though the length of the recordings limits the temporal resolution attainable.

## Conclusion

Here we have shown that low-dose ketamine triggers cardiovascular response reliably, i.e. it raises the heart rate. At the same time ketamine shifts the E:I balance, proxied by the aperiodic spectral exponent, robustly toward excitation in eyes-closed resting-state EEG data. A main contribution of our study lies in demonstrating that the cardiovascular response does not account for observed cortical E:I shift: controlling for heart rate left the ketamine effect not only intact but numerically stronger. Of practical relevance to a wider field of translational neuroscience and psychiatry, non-invasive proxies of E:I balance in neuroscience and psychiatry might thus remain even underestimated when not accounting for the systemic physiological response.

## Data availability statement

Data and code will be available on OSF.

## Acknowledgments

Hannah Schewe, Luisa Lindemann, Anne Herrmann, Ludmila Skrum, and Frederic Beba helped acquire the data.

## Author contributions

Conceptualization – ST (Sarah Tune), JO (Jonas Obleser), Alex Tzabazis (AT), Carla Nau (CN), Justus P. Student (JPS); Methodology – ST, JO, Judith Kunze (JK), JPS, Stefanie Schmidt (SS), Harald Ihmsen (HI), Henrik Oster (HO); Software – ST, JK; Formal analysis – ST, JPS, JO, JK; Investigation – ST, JPS, JK, AT, Benedikt Lorenz (BL); Resources – HO, SS, HI, JPS, CN; Data curation – JPS, JK; Writing – original draft – JPS, JO, JK; Writing – review & editing – JK, JPS, ST, HO, SS, HI, AT, BL, CN, JO; Visualization – ST, JPS, JK; Supervision – JO, CN, AT, BL; Project administration – ST, JO, JPS, CN; Funding acquisition — JO, CN, HO

