## Supplemental Materials for "Low-dose ketamine tilts the cortical excitation–inhibition balance irrespective of systemic physiological response"

### Supplementary Results

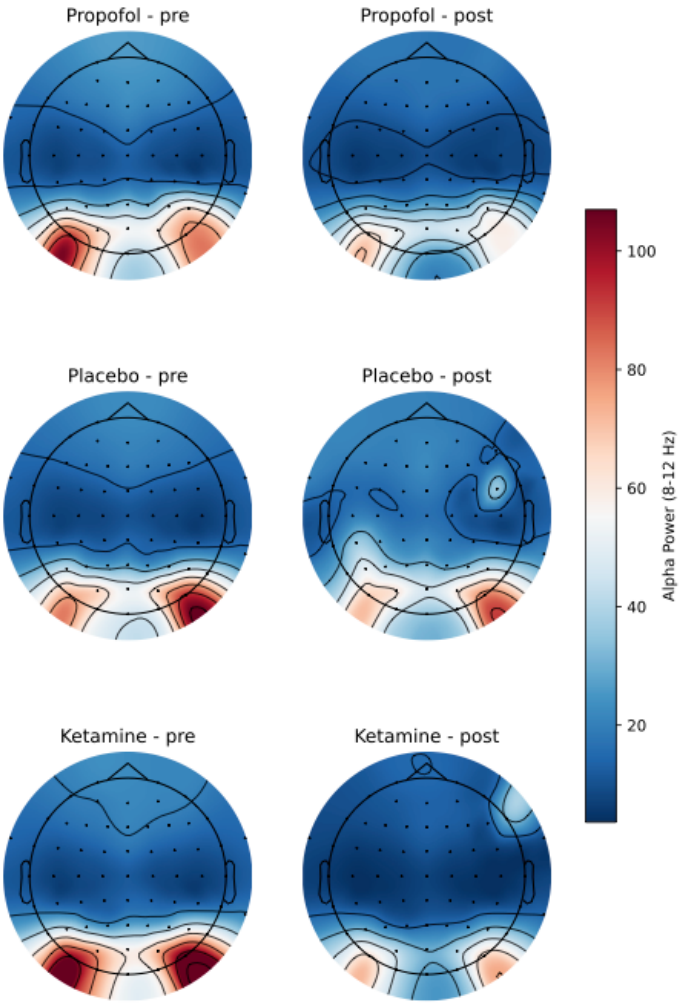


**Figure S1: Grand-average scalp topographies** of 1/f-corrected alpha power (8–12 Hz) at rest (eyes closed) before (left) and after (right) drug administration for propofol (top), placebo (middle), and ketamine (bottom). FOOOF models were fit per channel (see Methods); maps show the mean across N = 25 participants, plotted on a common colour scale.

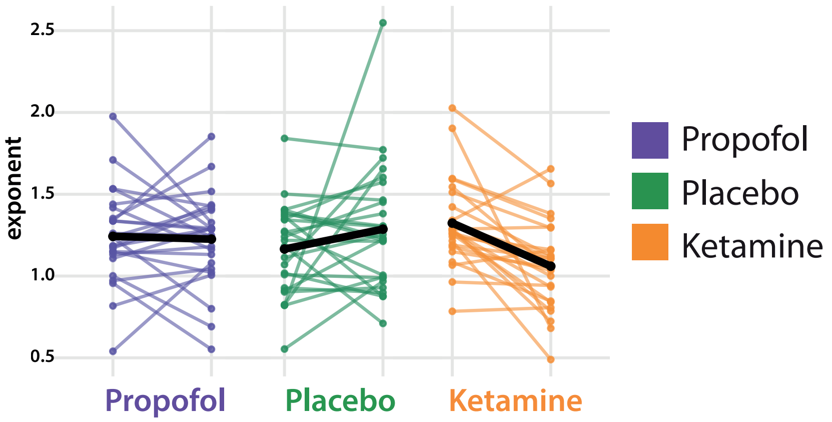


**Figure S2: Aperiodic exponent estimated alternatively over 1–50 Hz**, pre to post, per condition. Ketamine flattened the spectrum, i.e. the exponent decreased, whereas it increases under placebo. On average, the exponent stayed constant under propofol.

**Table S1: Subject characteristics.** Listed units are height in cm, weight in kg, and body mass index (BMI) in kg / m². Marked subjects completed all trials and were included in the final analyses.

| Subject | Age | Height | Weight | BMI | Sex | Completed |
|---|---|---|---|---|---|---|
| 001 | 20 | 172 | 60 | 20.3 | female | yes |
| 002 | 26 | 178 | 70 | 22.1 | female | yes |
| 003 | 20 | 164 | 58 | 21.6 | female | yes |
| 004 | 41 | 176 | 73 | 23.6 | female | yes |
| 005 | 20 | 172 | 70 | 23.7 | female | yes |
| 006 | 21 | 165 | 55 | 20.2 | female | no |
| 007 | 22 | 182 | 86 | 26.0 | male | yes |
| 008 | 25 | 178 | 76 | 24.0 | male | yes |
| 009 | 21 | 174 | 71 | 23.5 | female | yes |
| 010 | 21 | 172 | 71 | 24.0 | female | yes |
| 011 | 20 | 179 | 73 | 22.8 | male | yes |
| 012 | 22 | 170 | 70 | 24.2 | female | yes |
| 013 | 20 | 194 | 88 | 23.4 | male | yes |
| 014 | 22 | 183 | 67 | 20.0 | male | yes |
| 015 | 22 | 180 | 78 | 24.1 | female | yes |
| 016 | 20 | 173 | 64 | 21.4 | female | no |
| 017 | 22 | 196 | 90 | 23.4 | male | yes |
| 018 | 25 | 167 | 60 | 21.5 | female | yes |
| 019 | 31 | 178 | 70 | 22.1 | female | yes |
| 020 | 25 | 170 | 58 | 20.1 | female | no |
| 021 | 24 | 160 | 49 | 19.1 | female | no |
| 022 | 22 | 174 | 71 | 23.5 | male | no |

| | | | | | | |
|---|---|---|---|---|---|---|
| 023 | 23 | 180 | 72 | 22.2 | male | yes |
| 024 | 28 | 168 | 55 | 19.5 | female | yes |
| 025 | 21 | 160 | 45 | 17.6 | female | yes |
| 026 | 25 | 180 | 70 | 21.6 | male | yes |
| 027 | 23 | 165 | 57 | 20.9 | female | yes |
| 028 | 26 | 165 | 57 | 20.9 | female | yes |
| 029 | 19 | 196 | 85 | 22.1 | male | yes |
| 030 | 22 | 185 | 75 | 21.9 | male | no |
| 031 | 24 | 157 | 49 | 19.9 | female | yes |
| 032 | 28 | 182 | 93 | 28.1 | male | no |

**Table S2**

*Linear mixed-effects model predicting post-infusion aperiodic exponent*

| Predictor | *b* | *SE* | 95% CI | *df* | *t* | *p* |
|---|---|---|---|---|---|---|
| *Fixed effects* | | | | | | |
| Intercept | 0.00 | 0.13 | [−0.27, 0.27] | 18.28 | 0.00 | > .999 |
| Baseline exponent (*z*) | 0.16 | 0.12 | [−0.08, 0.39] | 59.75 | 1.35 | .183 |
| Ketamine vs. placebo | −0.75 | 0.24 | [−1.22, −0.27] | 44.80 | −3.17 | .003 |
| Propofol vs. placebo | −0.25 | 0.23 | [−0.71, 0.21] | 41.57 | −1.09 | .282 |
| Session (*z*) | 0.23 | 0.09 | [0.04, 0.42] | 41.27 | 2.47 | .018 |
| *Random effects* | *Variance* | *SD* | | | | |
| Participant (intercept) | 0.203 | 0.450 | | | | |
| Residual | 0.639 | 0.800 | | | | |

*Note.* *N* = 75 observations from 25 participants. Condition was entered with simple contrasts (contr.treatment(3) – 1/3) and placebo as the reference level; each condition coefficient therefore estimates the difference between that drug and placebo, and the intercept estimates the unweighted grand mean. Degrees of freedom and *p*-values were obtained with Satterthwaite's approximation; confidence intervals are Wald intervals. All z-standardized variables are expressed in standard deviation units. Intraclass correlation = .24. CI = confidence interval.

**Table S2.1**

*Type III analysis of variance for the model predicting post-infusion aperiodic exponent*

| Effect | *SS* | *MS* | $df_{num}$ | $df_{den}$ | *F* | *p* |
|---|---|---|---|---|---|---|
| Baseline exponent (*z*) | 1.16 | 1.16 | 1 | 59.75 | 1.81 | .183 |
| Condition | 6.68 | 3.34 | 2 | 42.66 | 5.22 | .009 |
| Session (*z*) | 3.91 | 3.91 | 1 | 41.27 | 6.11 | .018 |

*Note.* *N* = 75 observations from 25 participants. Type III sums of squares with Satterthwaite's approximation.

**Table 3**

*Linear mixed-effects model predicting post-infusion heart rate*

| Predictor | *b* | *SE* | 95% CI | *df* | *t* | *p* |
|---|---|---|---|---|---|---|
| *Fixed effects* | | | | | | |
| Intercept | 0.00 | 0.12 | [−0.25, 0.25] | 17.69 | 0.00 | > .999 |
| Baseline heart rate (*z*) | 0.40 | 0.08 | [0.25, 0.56] | 67.56 | 5.25 | < .001 |
| Ketamine vs. placebo | 0.79 | 0.13 | [0.52, 1.06] | 41.15 | 5.92 | < .001 |
| Propofol vs. placebo | −0.13 | 0.13 | [−0.40, 0.14] | 41.05 | −0.99 | .325 |
| *Random effects* | *Variance* | *SD* | | | | |
| Participant (intercept) | 0.288 | 0.536 | | | | |
| Residual | 0.216 | 0.465 | | | | |

*Note.* *N* = 75 observations from 25 participants. Condition was entered with simple contrasts (contr.treatment(3) − 1/3) and placebo as the reference level; each condition coefficient therefore estimates the difference between that drug and placebo, and the intercept estimates the unweighted grand mean. Degrees of freedom and *p*-values were obtained with Satterthwaite's approximation; confidence intervals are Wald intervals. All z-standardized variables are expressed in standard deviation units. Intraclass correlation = .57. CI = confidence interval.

**Table S3.1**

*Type III analysis of variance for the model predicting post-infusion heart rate*

| Effect | *SS* | *MS* | $df_{num}$ | $df_{den}$ | *F* | *p* |
|---|---|---|---|---|---|---|
| Baseline heart rate (*z*) | 5.97 | 5.97 | 1 | 67.56 | 27.61 | < .001 |
| Condition | 12.38 | 6.19 | 2 | 40.83 | 28.66 | < .001 |

*Note.* *N* = 75 observations from 25 participants. Type III sums of squares with Satterthwaite's approximation.

**Table S4**

*Linear mixed-effects model predicting post-infusion mean arterial pressure*

| Predictor | *b* | *SE* | 95% CI | *df* | *t* | *p* |
|---|---|---|---|---|---|---|
| *Fixed effects* | | | | | | |
| Intercept | −0.02 | 0.07 | [−0.17, 0.12] | 16.92 | −0.34 | .738 |
| Baseline MAP (*z*) | 0.55 | 0.06 | [0.42, 0.67] | 48.88 | 8.68 | < .001 |
| Ketamine vs. placebo | 1.36 | 0.12 | [1.12, 1.61] | 34.25 | 11.42 | < .001 |
| Propofol vs. placebo | −0.36 | 0.12 | [−0.61, −0.11] | 34.66 | −2.93 | .006 |
| *Random effects* | *Variance* | *SD* | | | | |
| Participant (intercept) | 0.062 | 0.249 | | | | |
| Residual | 0.158 | 0.398 | | | | |

*Note.* $N$ = 66 observations from 25 participants. Condition was entered with simple contrasts (contr.treatment(3) − 1/3) and placebo as the reference level; each condition coefficient therefore estimates the difference between that drug and placebo, and the intercept estimates the unweighted grand mean. Degrees of freedom and $p$-values were obtained with Satterthwaite's approximation; confidence intervals are Wald intervals. All z-standardized variables are expressed in standard deviation units. Intraclass correlation = .28. MAP = mean arterial pressure; CI = confidence interval.

**Table S4.1**

*Type III analysis of variance for the model predicting post-infusion mean arterial pressure*

| Effect | *SS* | *MS* | $df_{num}$ | $df_{den}$ | *F* | *p* |
|---|---|---|---|---|---|---|
| Baseline MAP (*z*) | 11.94 | 11.94 | 1 | 48.88 | 75.41 | < .001 |
| Condition | 34.89 | 17.45 | 2 | 34.86 | 110.21 | < .001 |

*Note.* *N* = 66 observations from 25 participants. Type III sums of squares with Satterthwaite's approximation. MAP = mean arterial pressure.

**Table S5**

*Linear mixed-effects model predicting post-infusion alpha power*

| Predictor | *b* | *SE* | 95% CI | *df* | *t* | *p* |
|---|---|---|---|---|---|---|
| *Fixed effects* | | | | | | |
| Intercept | −0.03 | 0.07 | [−0.17, 0.12] | 25.05 | −0.39 | .697 |
| Baseline alpha power (*z*) | 0.85 | 0.07 | [0.72, 0.99] | 34.10 | 12.71 | < .001 |
| Ketamine vs. placebo | −0.51 | 0.12 | [−0.75, −0.26] | 42.30 | −4.10 | < .001 |
| Propofol vs. placebo | −0.25 | 0.12 | [−0.50, 0.00] | 42.41 | −2.06 | .046 |
| Session (*z*) | −0.11 | 0.05 | [−0.21, 0.00] | 43 | −2.09 | .043 |
| *Random effects* | *Variance* | *SD* | | | | |
| Participant (intercept) | 0.060 | 0.245 | | | | |
| Residual | 0.159 | 0.398 | | | | |

*Note. N* = 65 observations from 25 participants. Condition was entered with simple contrasts (contr.treatment(3) − 1/3) and placebo as the reference level; each condition coefficient therefore estimates the difference between that drug and placebo, and the intercept estimates the unweighted grand mean. Degrees of freedom and *p*-values were obtained with Satterthwaite's approximation; confidence intervals are Wald intervals. All z-standardized variables are expressed in standard deviation units. Intraclass correlation = .27. CI = confidence interval.

**Table S5.1**

*Type III analysis of variance for the model predicting post-infusion alpha power*

| Effect | *SS* | *MS* | $df_{num}$ | $df_{den}$ | *F* | *p* |
|---|---|---|---|---|---|---|
| Baseline alpha power (*z*) | 25.61 | 25.61 | 1 | 34.10 | 161.59 | < .001 |
| Condition | 2.66 | 1.33 | 2 | 42.49 | 8.41 | < .001 |
| Session (*z*) | 0.69 | 0.69 | 1 | 43.00 | 4.35 | .043 |

*Note.* *N* = 65 observations from 25 participants. Type III sums of squares with Satterthwaite's approximation.

**Table S6**

*Linear mixed-effects model predicting post-infusion aperiodic exponent (30–50 Hz)*

| Predictor | *b* | *SE* | 95% CI | *df* | *t* | *p* |
|---|---|---|---|---|---|---|
| *Fixed effects* | | | | | | |
| Intercept | 0.00 | 0.10 | [−0.19, 0.19] | 70 | 0.00 | > .999 |
| Baseline exponent (*z*) | 0.45 | 0.10 | [0.25, 0.65] | 70 | 4.42 | < .001 |
| Ketamine vs. placebo | −0.62 | 0.24 | [−1.11, −0.14] | 70 | −2.55 | .013 |
| Propofol vs. placebo | 0.40 | 0.24 | [−0.08, 0.88] | 70 | 1.68 | .097 |
| Session (*z*) | 0.00 | 0.10 | [−0.20, 0.20] | 70 | −0.01 | .992 |
| *Random effects* | *Variance* | *SD* | | | | |
| Participant (intercept) | 0.000 | 0.000 | | | | |
| Residual | 0.703 | 0.839 | | | | |

*Note.* $N = 75$ observations from 25 participants. Condition was entered with simple contrasts (contr.treatment(3) − 1/3) and placebo as the reference level; each condition coefficient therefore estimates the difference between that drug and placebo, and the intercept estimates the unweighted grand mean. The random-intercept variance was estimated at the boundary (zero), that is, the fit was singular; the fixed-effects estimates are therefore equivalent to those of an ordinary least-squares model. Degrees of freedom and *p*-values were obtained with Satterthwaite's approximation; confidence intervals are Wald intervals. All z-standardized variables are expressed in standard deviation units. Intraclass correlation = .00. CI = confidence interval.

**Table S6.1**

*Type III analysis of variance for the model predicting post-infusion aperiodic exponent (30–50 Hz)*

| Effect | *SS* | *MS* | $df_{\text{num}}$ | $df_{\text{den}}$ | *F* | *p* |
|---|---|---|---|---|---|---|
| Baseline exponent (*z*) | 13.71 | 13.71 | 1 | 70 | 19.51 | < .001 |
| Condition | 13.05 | 6.53 | 2 | 70 | 9.28 | < .001 |
| Session (*z*) | 0.00 | 0.00 | 1 | 70 | 0.00 | .992 |

*Note.* *N* = 75 observations from 25 participants. Type III sums of squares with Satterthwaite's approximation. The model fit was singular.

**Table S7**

*Linear mixed-effects model predicting post-infusion aperiodic exponent (30–50 Hz), controlling for heart rate*

| Predictor | *b* | *SE* | 95% CI | *df* | *t* | *p* |
|---|---|---|---|---|---|---|
| *Fixed effects* | | | | | | |
| Intercept | 0.01 | 0.10 | [−0.18, 0.20] | 69 | 0.07 | .941 |
| Baseline exponent (*z*) | 0.46 | 0.10 | [0.26, 0.66] | 69 | 4.57 | < .001 |
| Ketamine vs. placebo | −0.75 | 0.26 | [−1.26, −0.24] | 69 | −2.93 | .005 |
| Propofol vs. placebo | 0.44 | 0.24 | [−0.03, 0.92] | 69 | 1.86 | .067 |
| Heart rate (*z*) | 0.17 | 0.11 | [−0.05, 0.40] | 69 | 1.55 | .126 |
| Session (*z*) | 0.00 | 0.10 | [−0.19, 0.20] | 69 | 0.03 | .976 |
| *Random effects* | *Variance* | *SD* | | | | |
| Participant (intercept) | 0.000 | 0.000 | | | | |
| Residual | 0.689 | 0.830 | | | | |

*Note.* *N* = 75 observations from 25 participants. Condition was entered with simple contrasts (contr.treatment(3) − 1/3) and placebo as the reference level; each condition coefficient therefore estimates the difference between that drug and placebo, and the intercept estimates the unweighted grand mean. The random-intercept variance was estimated at the boundary (zero), that is, the fit was singular; the fixed-effects estimates are therefore equivalent to those of an ordinary least-squares model. Degrees of freedom and *p*-values were obtained with Satterthwaite's approximation; confidence intervals are Wald intervals. All z-standardized variables are expressed in standard deviation units. Intraclass correlation = .00. CI = confidence interval.

**Table S7.1**

*Type III analysis of variance for the model predicting post-infusion aperiodic exponent (30–50 Hz), controlling for heart rate*

| Effect | *SS* | *MS* | $df_{num}$ | $df_{den}$ | *F* | *p* |
|---|---|---|---|---|---|---|
| Baseline exponent (*z*) | 14.36 | 14.36 | 1 | 69 | 20.84 | < .001 |
| Condition | 14.57 | 7.29 | 2 | 69 | 10.57 | < .001 |
| Heart rate (*z*) | 1.66 | 1.66 | 1 | 69 | 2.40 | .126 |
| Session (*z*) | 0.00 | 0.00 | 1 | 69 | 0.00 | .976 |

*Note.* *N* = 75 observations from 25 participants. Type III sums of squares with Satterthwaite’s approximation. The model fit was singular.

**Table S8**

*Linear mixed-effects model predicting post-infusion dissociation*

| Predictor | *b* | *SE* | 95% CI | *df* | *t* | *p* |
|---|---|---|---|---|---|---|
| *Fixed effects* | | | | | | |
| Intercept | −0.52 | 0.33 | [−1.19, 0.15] | 39.32 | −1.58 | .123 |
| Baseline CADSS (*z*) | 0.07 | 0.10 | [−0.14, 0.28] | 45.90 | 0.68 | .502 |
| Ketamine vs. placebo | 1.53 | 0.23 | [1.04, 2.01] | 23.91 | 6.51 | < .001 |
| Session | −0.02 | 0.14 | [−0.31, 0.27] | 34.91 | −0.17 | .866 |
| *Random effects* | *Variance* | *SD* | | | | |
| Participant (intercept) | 0.052 | 0.228 | | | | |
| Residual | 0.658 | 0.811 | | | | |

*Note.* *N* = 50 observations from 25 participants. Condition was dummy-coded with placebo as the reference level (only placebo and ketamine sessions contributed to this model); the intercept therefore estimates the expected value in the placebo condition when all covariates are at zero. Degrees of freedom and *p*-values were obtained with Satterthwaite's approximation; confidence intervals are Wald intervals. All z-standardized variables are expressed in standard deviation units. Intraclass correlation = .07. CADSS = Clinician-Administered Dissociative States Scale; CI = confidence interval.

**Table S9**

*Linear mixed-effects model predicting post-infusion dissociation, controlling for heart rate*

| Predictor | *b* | *SE* | 95% CI | *df* | *t* | *p* |
|---|---|---|---|---|---|---|
| *Fixed effects* | | | | | | |
| Intercept | –0.51 | 0.33 | [–1.18, 0.16] | 39.02 | –1.53 | .133 |
| Baseline CADSS (*z*) | 0.06 | 0.10 | [–0.15, 0.27] | 45 | 0.57 | .569 |
| Ketamine vs. placebo | 1.67 | 0.26 | [1.14, 2.21] | 29.98 | 6.44 | < .001 |
| Heart rate (*z*) | –0.20 | 0.14 | [–0.49, 0.08] | 29.77 | –1.45 | .158 |
| Session | –0.05 | 0.15 | [–0.34, 0.24] | 35.22 | –0.35 | .731 |
| *Random effects* | *Variance* | *SD* | | | | |
| Participant (intercept) | 0.008 | 0.091 | | | | |
| Residual | 0.685 | 0.828 | | | | |

*Note.* $N = 50$ observations from 25 participants. Condition was dummy-coded with placebo as the reference level (only placebo and ketamine sessions contributed to this model); the intercept therefore estimates the expected value in the placebo condition when all covariates are at zero. Degrees of freedom and *p* values were obtained with Satterthwaite's approximation; confidence intervals are Wald intervals. All z-standardized variables are expressed in standard deviation units. Intraclass correlation = .01. CADSS = Clinician-Administered Dissociative States Scale; CI = confidence interval.

**Table S10**

*Linear mixed-effects model predicting post-infusion dissociation, controlling for the aperiodic exponent*

| Predictor | *b* | *SE* | 95% CI | *df* | *t* | *p* |
|---|---|---|---|---|---|---|
| *Fixed effects* | | | | | | |
| Intercept | −0.52 | 0.33 | [−1.20, 0.15] | 38.55 | −1.56 | .126 |
| Baseline CADSS (*z*) | 0.07 | 0.11 | [−0.14, 0.28] | 44.93 | 0.65 | .520 |
| Ketamine vs. placebo | 1.49 | 0.24 | [0.99, 1.99] | 24.47 | 6.12 | < .001 |
| Aperiodic exponent (*z*) | −0.11 | 0.14 | [−0.39, 0.17] | 44.36 | −0.79 | .433 |
| Session | −0.03 | 0.14 | [−0.32, 0.27] | 34.42 | −0.19 | .848 |
| *Random effects* | *Variance* | *SD* | | | | |
| Participant (intercept) | 0.036 | 0.189 | | | | |
| Residual | 0.679 | 0.824 | | | | |

*Note.* $N$ = 50 observations from 25 participants. Condition was dummy-coded with placebo as the reference level (only placebo and ketamine sessions contributed to this model); the intercept therefore estimates the expected value in the placebo condition when all covariates are at zero. Degrees of freedom and *p*-values were obtained with Satterthwaite's approximation; confidence intervals are Wald intervals. All z-standardized variables are expressed in standard deviation units. Intraclass correlation = .05. CADSS = Clinician-Administered Dissociative States Scale; CI = confidence interval.

**Table S11**

*Linear mixed-effects model predicting post-infusion dissociation, controlling for heart rate and the aperiodic exponent*

| Predictor | *b* | *SE* | 95% CI | *df* | *t* | *p* |
|---|---|---|---|---|---|---|
| *Fixed effects* | | | | | | |
| Intercept | −0.51 | 0.33 | [−1.18, 0.17] | 44 | −1.52 | .136 |
| Baseline CADSS (*z*) | 0.06 | 0.10 | [−0.15, 0.27] | 44 | 0.55 | .588 |
| Ketamine vs. placebo | 1.64 | 0.27 | [1.09, 2.18] | 44 | 6.06 | < .001 |
| Aperiodic exponent (*z*) | −0.09 | 0.14 | [−0.37, 0.19] | 44 | −0.64 | .523 |
| Heart rate (*z*) | −0.19 | 0.14 | [−0.48, 0.09] | 44 | −1.36 | .182 |
| Session | −0.05 | 0.15 | [−0.35, 0.24] | 44 | −0.36 | .720 |
| *Random effects* | *Variance* | *SD* | | | | |
| Participant (intercept) | 0.000 | 0.000 | | | | |
| Residual | 0.703 | 0.838 | | | | |

*Note.* $N$ = 50 observations from 25 participants. Condition was dummy-coded with placebo as the reference level (only placebo and ketamine sessions contributed to this model); the intercept therefore estimates the expected value in the placebo condition when all covariates are at zero. The random-intercept variance was estimated at the boundary (zero), that is, the fit was singular; the fixed-effects estimates are therefore equivalent to those of an ordinary least-squares model. Degrees of freedom and *p*-values were obtained with Satterthwaite's approximation; confidence intervals are Wald intervals. All z-standardized variables are expressed in standard deviation units. Intraclass correlation = .00. CADSS = Clinician-Administered Dissociative States Scale; CI = confidence interval.
